# Basal forebrain and neural correlates of self-regulation traits in sustained attention

**DOI:** 10.1101/2025.08.05.668456

**Authors:** Chiara Orsini, Daniel A. Huber, Karin Labek, Julia E. Bosch, Roberto Viviani

## Abstract

Self-regulation is a human trait consistently associated with success in both academic and professional settings and to better mental health. Based on previous findings, we used functional imaging data in a sustained attention task to test three hypotheses on neural substrates associated with individual differences in self-regulation. The first linked higher self-regulation and cognitive control, predicting modulation of recruitment of prefrontal substrates. The second, originating in the animal literature, suggests increased recruitment of cholinergic substrates in the basal forebrain. The third predicted higher modulation of reward-sensitive regions in the brainstem in less regulated individuals for differences in reward levels during the task. The second hypothesis was confirmed by our study, which also provided suggestive evidence for the third hypothesis. Our data suggest that one mechanism of higher self-regulation in humans may ensue from greater activity in the cholinergic system to sustain attention during a cognitively simple task.

## 1 Introduction

Self-regulation is a predictor of health, well-being, and adaptive behavior (Bub et al., 2016; Congdon & Canli, 2005; Duckworth & Carlson, 2013; Moffitt et al., 2011; Pandey et al., 2018; Weidner et al., 2016). In psychology, self-regulation is an umbrella term that generally subtends the ability to utilize self-control when this is appropriate to attain a goal through one’s own cognition, emotions, and behavior (Baumeister et al., 2002; Bentivegna et al., 2024; Diamond, 2013; Matthews et al., 2000). Self-control is considered a component of inhibitory control and is the opposite of the tendency to be impulsive (see Table 1 for a clarification of the terminology used in the present paper); it includes the ability to stay on a task, opting not to disengage from it notwithstanding distractions (Diamond, 2013), and to concentrate (Baumeister et al., 2002). This theorization is also supported by recent behavioral evidence that found a positive association between sustained attention performance and trait self-regulation (Harwood et al., 2023; Mouloua & Shaw, 2023). The present study contributes to this field of research by investigating the neural correlates of individual differences in self-regulation using functional neuroimaging in a sustained attention task (Orsini et al., 2026).

**Table 1.** Self-regulation terminology as used in this manuscript.

| Term | Meaning |
| --- | --- |
| self-regulation | Ability to organize and control one's psychological processes to achieve a goal, as when persisting in pursuing this goal also in the absence of immediate rewards. It is adaptive. |
| self-control | Ability to control one's impulses. It is a requirement to achieve self-regulation, but unlike this latter may also be applied in an excessive or inappropriate manner. |
| cognitive control | The aspect of self-control that depends on/consists of cognitive capacity, such as the recruitment of attentional processes or executive functions. |
| sustained attention | The endogenous allocation of attentional resources to stay on task and mitigate goal-neglect when continuously responding to repetitive stimuli (Coull, 1998; Robertson & O'Connell, 2010). Evidence supports its involvement after a few seconds of work (Langner & Eickhoff, 2013; Orsini et al., 2026; Robertson & O'Connell, 2010). |

A considerable amount of empirical evidence about the positive outcomes associated with self-regulation has been obtained with scales such as the Self-Control Scale (SCS), which has been shown to correlate with better achievements and task performance, better impulse control (for example, in eating behavior), positive psychological adjustment and self-esteem, better interpersonal relations, taking responsibility for transgressions and a moral emotional style (Tangney et al., 2004). Over the years, however, several other approaches have been proposed to empirically assess self-regulation and characterize self-control operationally.

One such characterization has been the goal-oriented ability to delay a gratification or the acquisition of a benefit to achieve a better outcome at a later point (Mischel et al., 1989). In this classic literature, Mischel and colleagues extensively reported the presence of an association between self-control and attentional processes (Mischel & Ebbesen, 1970; Mischel et al., 1972; Mischel et al., 1989). More recently, Posner and colleagues bolstered this hypothesis and argued that high-order attentional processes, especially those of executive nature, drive self-control (cognitive control, Posner & Rothbart, 2000; Posner et al., 2013; Rueda et al., 2004, 2005). From a neural perspective, Posner et al. (2007) argued that self-control mechanisms are likely localized in the executive attention network and particularly in the anterior cingulate cortex (ACC)/midfrontal cortex, a view substantiated by neuroimaging evidence (Banfield et al., 2004). According to an influential model, ACC is also involved in computing the trade-off between the benefits of allocating cognition and the effort that goes with the application of cognition (Shenhav et al., 2016; Shenhav et al., 2017). A related approach has proposed to assess self-control as the capacity to inhibit prepotent responses, as in go-no go or Stroop tasks (Casey et al., 2011; Chamberlain & Sahakian, 2007), similarly identifying its correlates in areas of the prefrontal cortex such as the ACC or the inferior frontal gyrus (IFG).

A complementary perspective has been provided by the animal literature. In this literature, the neurotransmitter acetylcholine (ACh) in the basal forebrain (BF) mediates cognitive effort, and ACh cortical efflux increases in the case of cognitive challenges and counters attentional decrements (Sarter et al., 2006). Animal studies have also provided evidence of the effects of ACh agonists/antagonists on effort, attention, and on individual differences in performance (Hosking et al., 2014), highlighting the connection between the cholinergic system and individual phenotypes.

Reward has also been repeatedly shown to facilitate sustained attention and cognitive effort more generally, increasing attentional performance (Dutra et al., 2018; Esterman et al., 2017; Esterman et al., 2014; Massar et al., 2016; Tomporowski & Tinsley, 1996). By converse, sensitivity to rewards characterizes impulsivity (Evenden, 1999), consistent with the findings in delay discounting studies (Göllner et al., 2018; Mischel et al., 1989). Neuroimaging studies have reported reduced activation in dopaminergic regions in impulsive individuals resisting pursuit of immediate rewards in favor of a long-term goal (Diekhof et al., 2012).

Sustained attention (Table 1) is the cognitive process involved in remaining task-focused over time (Coull, 1998), often investigated empirically in the context of long repetitive attentional tasks. However, a form of sustained attention is already recruited during a short on-task period (Posner & Boies, 1971) of 10 seconds or less (Langner & Eickhoff, 2013). At this short time on task, a cognitive process may be required to compensate momentary lapses of goal-directed attention (Langner & Eickhoff, 2013; Robertson & O’Connell, 2010). This distraction-preventing function may differ from compensating the mental fatigue and the vigilance decrements that are typically associated with long times on task (for a detailed discussion see Orsini et al., 2026). This form of sustained attention is required also in the absence of working memory requirements in the task (as in the tasks traditionally used to investigate sustained attention, see Mackworth, 1948). By focusing on executive function as the primary source of cognitive control, the functional imaging literature has left unexplored the possible existence of individual differences in self-regulation in the substrates recruited by sustained attention.

In the sustained attention task of the present study, participants responded to the appearance of a stimulus on the left or right side of the screen by pressing a button on the same side. These target stimuli were grouped in blocks of 12-13 lasting about 15 seconds, referred to as ‘foraging blocks’ (Orsini et al., 2026). Correct presses were rewarded with 1 or 20 virtual cents depending on the block in which the stimuli were presented. Blocks were preceded by the display of a cue that informed participants of the reward level of the block (Figure 1A). The total reward in real money could only be obtained if they responded correctly to enough stimuli, thus introducing a tension between the necessity to stay on task and the amount of reward collected at each trial (i.e., continuous performance was required even if the reward was low). The threshold to obtain the real money was reached only at the end of the task. Importantly, the time on task was modelled for the block, i.e., was reset at the beginning of each block (Figure 1B), avoiding confounders that arise at longer time scales.

**Figure 1.**
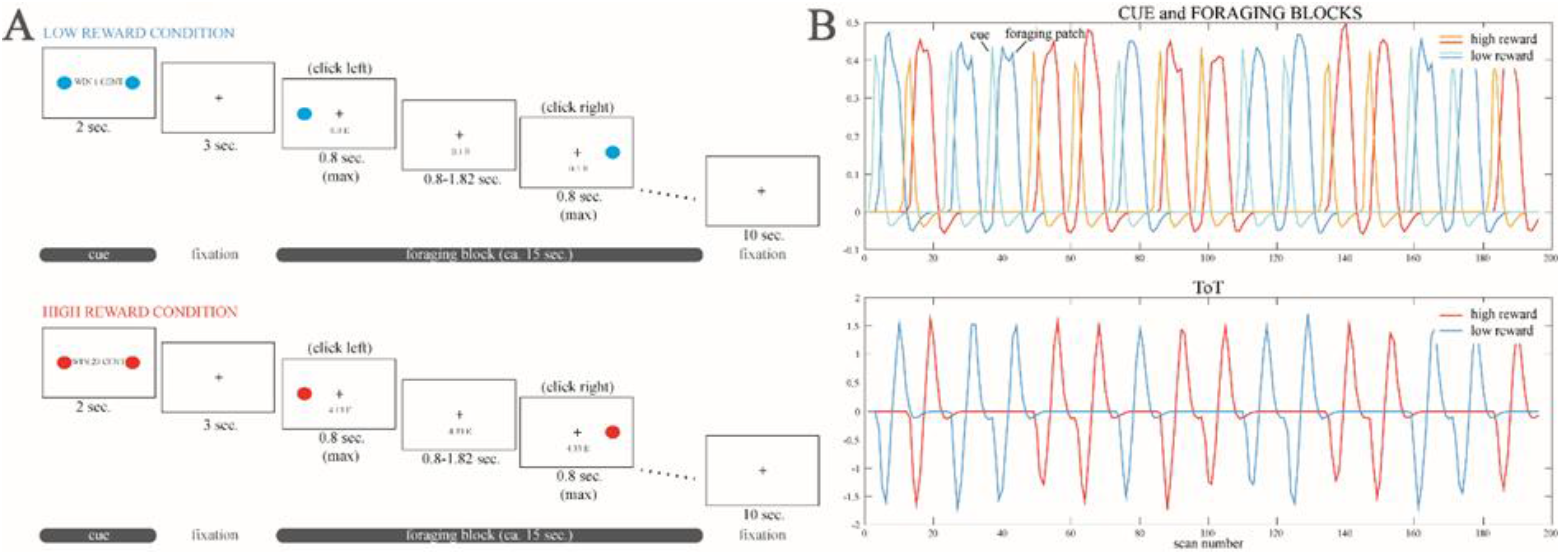
A: Schematic rendition of the task used in the study. Participants had to click on a right or left button depending on the side on which a dot appeared on the screen during blocks of trials in which reward for correct responses could be low or high. Each foraging block was preceded by a cue announcing the reward level of the following block. B: regressors used in the study. ToT: time on task.

In this paradigm, we have previously shown that the short time on task evoked activity in widespread cortical areas peaking in the ACC and right IFG (Orsini et al., 2026). These areas correspond to those that, in neuroimaging literature, have been shown to be active in sustained attention tasks (Langner & Eickhoff, 2013). Multiple subcortical substrates were also involved such as the ventral tegmental area/substantia nigra (VTA/SN), the nucleus accumbens (NAcc), and the cholinergic portion of the nucleus basalis of Meynert (NBM/Ch4), located in the BF (Orsini et al., 2026). The subcortical effects in VTA/SN and NBM/Ch4 were modulated by reward levels. These neural systems were engaged while maintaining behavioral uniformity across participants. This makes this paradigm a controlled probe of neural systems where neural differences reflect trait-related processing differences rather than downstream consequences of performance failure, such as error monitoring, frustration, or coping with misses. The short time on task also avoids the confounders of fatigue, learning effects, and scanner drift.

The data from this study, therefore, offered the opportunity to test the role of these neuroimaging phenotypes in individual differences in self-regulation, which was assessed with the SCS scale in a group of healthy participants (N=203). The first hypothesis we wanted to test is the possible differential recruitment of ACC and IFG in highly regulated individuals, as in the functional imaging literature about cognitive control (hypothesis 1). Based on the animal literature on the role of cholinergic systems in attentional performance, we hypothesized increased recruitment of the NBM/Ch4 in individuals with higher self-regulation (hypothesis 2). Finally, we hypothesized that changes in activity accompanying reward conditions would be more marked in less self-regulated individuals, which might be more sensitive to rewards (hypothesis 3). To our knowledge, none of these hypotheses has been tested in a sustained attention task; furthermore, nothing is known about the possible role of the BF in human self-regulation, as current evidence is provided by animal studies using what appear to be related constructs.

We conducted region of interest (ROI) analyses on ACC, IFG, Ch4, NAcc, and VTA/SN as well as whole-brain analyses to detect correlates in areas not covered by our hypotheses. The main novel finding of our study was that highly regulated individuals presented a greater involvement of the BF over the time on task, irrespective of reward levels, confirming the second of our hypotheses. This finding was also significant in the whole brain analyses, which did not show any other result surviving correction for multiple testing. Finally, the VTA/SN was involved in the effect of self-regulation in the interaction between time on task and reward, as in the third of our hypotheses, but this involvement failed to survive stringent levels of significance correction.

## 2 Materials and methods

### 2.1 Participants

The present study analyzed data from a previous genetic imaging research project (Orsini et al., 2026; Viviani, Messina, et al., 2020). Written informed consent was given by 441 healthy Caucasian volunteers, who were screened for psychiatric disorders. Exclusion criteria were addiction to alcohol or drugs, pregnancy or lactation, psychiatric or affective psychological disorders, severe acute or chronic diseases, metal implants, large tattoos, or tattoos near the head, and intake of psychoactive or long-term medications. Reasons for exclusion after data collection were clinical findings, artifacts, excessive movements, equipment failure, task administration or completion failures. Among the remaining 415 individuals, 203 individuals (114 females, age range 18-38, mean age 23.06 ± 3.88 years) completed the Self-Control Scale (Tangney et al., 2004) in its brief German version (Bertrams & Dickhäuser, 2009), and were therefore eligible for the present study. This scale was available for assessment in the second half of the study period and was given to all participants recruited in this half. Data collection of 20 participants was carried out at the German Center for Neurodegenerative Diseases (DZNE) located in Bonn, whereas the data of the remaining 183 participants were obtained at the University of Ulm.

The study (acronym: BrainCYP) was registered in the German Clinical Trials Register (DRKS-ID: 00011722). It was carried out in accordance with the Declaration of Helsinki guidelines and received approval from the Ethical Committees of both the University of Bonn (No. 33/15) and the University of Ulm (No. 01/15).

### 2.2 Experimental paradigm

The paradigm (Figure 1A) was described in Viviani, Dommes, et al. (2020). It consisted of a simple sustained attention task lasting about 8 minutes (Orsini et al., 2026; Viviani, Dommes, et al., 2020; Viviani, Messina, et al., 2020) where 12/13 trials were presented grouped within 16 ‘foraging blocks’. Every block lasted about 15s and consisted of the presentation of dots on the left or the right side of the screen, requiring participants to press a left or right button (Figure 1A). Before each block, a 2s cue was presented indicating the potential reward that could be earned during every trial within the upcoming block, which was 1 or 20 virtual cents for each correct response. According to a Poisson schedule, the interstimulus interval (ISI) varied from 0.80s to 1.82s (average value 1.23s). Because of the simplicity of the task, participants had a short time to detect the signal (maximum 0.8s). A 10s pause featuring the fixation cross followed each foraging phase before the next cue phase began. If 20 virtual euros or more were accumulated during the task, the earnings could be converted into real money. Prior to the session, participants were informed that staying on task, even in the low-reward trials, was essential to reaching this threshold, and they had the possibility to practice.

### 2.3 Rating scales

To evaluate individual differences in self-regulation, the short and German version of the Self-Control Scale (SCS-K-D) (Bertrams & Dickhäuser, 2009; Tangney et al., 2004) was administered. The statistical analyses were computed through the freely-available software R (R: The R Project for Statistical Computing, r-project.org, version 4.4.0) using the packages lme4 (Bates et al., 2015), lmerTest (Kuznetsova et al., 2017), dplyr (Wickham et al., 2023), and ggplot2 (Wickham, 2016) for plotting. To validate the scores of the SCS-K-D scale, we verified in our data that it could predict body-mass index (BMI) scores as reported in the literature (Cobb-Clark et al., 2023; Keller et al., 2016; Koike et al., 2016). In a linear model adjusted for acquisition site, age, and sex, self-regulation was a significant predictor of BMI (*t* = -2.51, *p* = 0.01). An increase of one point in the SCS-K-D corresponded to a decrease of 0.07 points in the BMI. Also, sex was associated with self-regulation scores, with females presenting higher scores in the questionnaire (*t* = 2.29, *p* = 0.02). The presence of an association between the SCS-K-D scores and behavioral performance (reaction times and accuracy) was also investigated. Briefly, reaction times were modeled with a linear mixed-effects model accounting for the effects of demographic (age, sex, acquisition site, SCS-K-D scores) and task-related variables (trial number within block, first trial, block number, switch, ISI, and reward level). This model included a random intercept for each subject. Accuracy was modeled with a binomial generalized linear mixed-effects model including the same predictors and a random intercept for each subject. For details, see the analysis in the Supplementary Materials.

### 2.4 fMRI data acquisition

Data were gathered at the German Center for Neurodegenerative Diseases (DZNE) located in Bonn and the Department of Psychiatry and Psychotherapy at the University of Ulm with a 3 T Siemens Skyra and a 3 T Siemens Prisma scanner, respectively. Both were fitted with 64-channel head coils. For the imaging protocol, a T2*-sensitive echo-planar imaging sequence was used (TR/TE = 2460/30ms; flip angle = 82°; FOV = 24cm; image matrix = 64 × 64 pixels; voxel size = 3 × 3mm). A total of 39 transverse slices, each 2.5mm thick, were acquired in ascending order. The gap between slices measured 0.5mm, giving an isotropic voxel size of 3mm. To account for the shorter T2* in high susceptibility and iron-rich regions, TE was gradually decreased by 8ms from slice 24 to slice 14, achieving a TE of 22ms for the first 14 slices at the bottom of the volume (for a description of the method, see Stöcker et al., 2006). Additionally, collected T1-weighted structural images were examined individually to evaluate the presence of abnormalities.

### 2.5 Data analysis

The present study analyzes data used in a previous genetic imaging study (Viviani, Messina, et al., 2020). Using the freely available software SPM12 (Welcome Trust Centre for Neuroimaging, http://www.fil.ion.ucl.ac.uk/spm/) running on MATLAB (The MathWorks, https://www.mathworks.com). Data were preprocessed with a realignment procedure, registered to the Montreal Neurological Institute (MNI) space, resampled to a voxel size of 1.5mm, and smoothed (Gaussian kernel 8mm FWHM). During registration, accurate realignment of subcortical structures was obtained by complementing the SPM spatial priors with one additional segment for the pallidum, the substantia nigra, and the red nucleus (see also Orsini et al., 2026; Viviani, Dommes, et al., 2020).

At the first level, cues and trials in block were separately modeled as zero-duration onset series, one per each rewarding level (for a total of four series, two for the cues and two for the trials in block), and convolved with the canonical hemodynamic response function. To model the effect of the time spent on task, a parametric modulator equal to the centered number of trials within a block was used. The parametric modulator coefficient represented the increase of brain metabolic activity within the block (see details in Orsini et al., 2026). The SPM default high-pass filter of 128s was applied. Data were denoised using the mean signal of the cranial bone (Huber et al., 2024), of the white matter, and of the cerebrospinal fluid as three separate confounding covariates in the first level model.

As in Orsini et al. (2026), two contrasts were brought to the second level: the time on task contrast, representing the brain activity variations over time within the blocks (time on task of about 15s), and the interaction between time on task and the rewarding context (high vs low). SCS-K-D individual scores were used as main regressor in the neuroimaging analysis to identify the patterns of activation associated with high or low trait self-regulation levels. In this model, acquisition site and participants’ sex and age were used as confounding covariates. Significance was assessed with permutation tests with *a priori-*defined regions of interest (ROI) and for the whole brain. ROI-based permutation tests were computed for ACC, IFG, Ch4, NAcc, and VTA/SN (8000 resamples, cluster-defining threshold *p* = 0.01, uncorrected), each with a mask of the corresponding region (see the Masks section below). Whole brain permutation tests (8000 resamples, cluster-defining threshold *p* = 0.001, uncorrected) were computed after applying a mask excluding the cerebellum and non-neuronal substrates (white matter and cerebrospinal fluid). The final results were visualized through the freely available software MRICron (Chris Rorden, https://people.cas.sc.edu/rorden/mricron). Brain regions were identified with the online tool available at http://www.talairach.org/ after conversion to the Talairach coordinates. This labelling was supplemented by the EBRAINS Multilevel Human Atlas (https://www.ebrains.eu/) for subcortical regions and by the MNI2Tal tool by BioImage Suite Web (https://bioimagesuiteweb.github.io/bisweb-manual/tools/mni2tal.html) for missing information about Brodmann areas.

### 2.6 Masks

The ACC and IFG masks were extracted from the data available at www.brain-development.org (Gousias et al., 2008; Hammers et al., 2003). The Ch4 mask was obtained from the SPM Anatomy Toolbox (Eickhoff et al., 2006; Eickhoff et al., 2007; Eickhoff et al., 2005; Zaborszky et al., 2008). The NAcc mask was retrieved from the Harvard Oxford Atlas (Desikan et al., 2006; Frazier et al., 2005; Goldstein et al., 2007; Makris et al., 2006), distributed by the FMRIB Software Library (https://fsl.fmrib.ox.ac.uk/). The VTA/SN mask was extracted from data available at https://www.adcocklab.org/neuroimaging-tools (Ballard et al., 2011; Murty et al., 2014). The ACC, IFG, and VTA/SN structural masks were finally combined with a functional mask of the simple effects of the two main contrasts (from the model without individual differences, Viviani, 2010), generating a combined functional/structural mask per substrate and contrast of interest.

## 3 Results

The present study focuses on the presence of functional brain phenotypes of trait self-regulation during the execution of a sustained attention task. The effects of time on task and its interaction with reward, regardless of individual differences, were reported by Orsini et al. (2026). Briefly, in that work, very small but significant performance decrements during the short time on task (both in terms of increased reaction times and hit rates) were reported. In the neuroimaging analyses, time on task and the interaction with reward levels were characterized by augmented activity in widespread cortical substrates (peaking in frontal areas in the ACC and IFG) and in subcortical regions such as the basal ganglia, BF, and VTA/SN. These findings were used together with anatomical information to define the *a priori* ROI of the analyses to test the three models of self-regulation mentioned in the Introduction (see Materials and methods for details).

### 3.1 Behavioral analysis

There were no significant differences in performance decrements associated with individual differences in self-regulation. SCS-K-D scores did not predict reaction times (*t* = 0.85, *p* = 0.40) and accuracy (*z* = -0.65, *p* = 0.52). For further details about the models, see the Supplementary Materials.

### 3.2 Association of self-regulation with time on task (hypotheses 1 and 2)

In the ROI analyses of time on task, self-regulation was associated with differences in activity in bilateral Ch4 (x, y, z: -17, -1, -12, *t* = 3.900, *p* = 0.001; and 20, 1, -10, *t* = 3.607, *p* = 0.001) and NAcc (-12, 7, -7, *t* = 3.930, *p* = 0.012; and 15, 16, -7, *t* = 3.393, *p* = 0.015; all significance levels at cluster level correction; for details and other corrections, see Table 1A in the Appendix). Larger increases in activity during time on task were associated with higher levels of self-regulation (Figure 2A and 2B). In contrast, no significant modulation of time on task activity associated with self-regulation levels were detected in the prefrontal ROIs.

**Figure 2.**
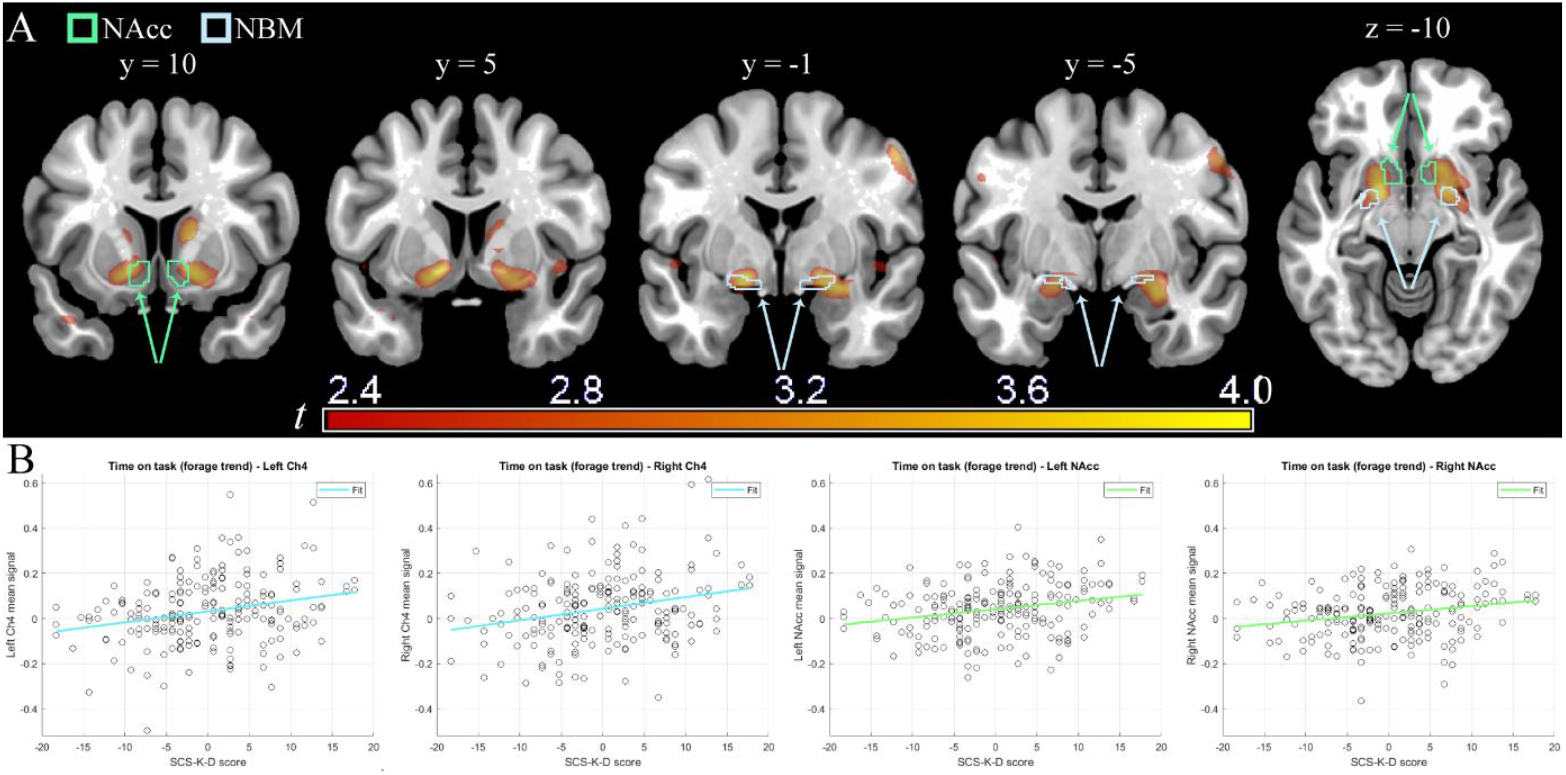
A: Whole brain functional activity in the interaction time on task × self-regulation scores (thresholded for illustration purposes at 2.35 < *t* < 4.00, *p* < 0.01 uncorrected). In outline, the ROI masks from the literature for the NAcc (in green) and NBM (in pale blue) are shown. B: Illustrative plots of participants’ Ch4 (pale blue) or NAcc (green) ROI mean signal extracted from the interaction time on task × self-regulation scores (y-axis) and SCS-K-D score (x-axis; see text for inference).

The whole brain analysis (Table 2A, Appendix) revealed that the modulation of activity in the NAcc associated with self-regulation levels was part of a larger and significant right-lateralized cluster centered on Ch4 and the adjacent ventral pallidum (x, y, z: 20, 1, -10, *t* = 3.607, *p* = 0.040), also extending towards the amygdala (x, y, z: 27, -4, -16, *t* = 3.891, *p* = 0.040). Involvement of the NAcc, while significant, was partial. A second cluster, lateralized on the left and similarly extending from the Ch4/ventral pallidum to partial NAcc components and central amygdala, reached trend-level significance (*p* = 0.079). No other region was significant in the whole brain analysis.

**Table 2A.**
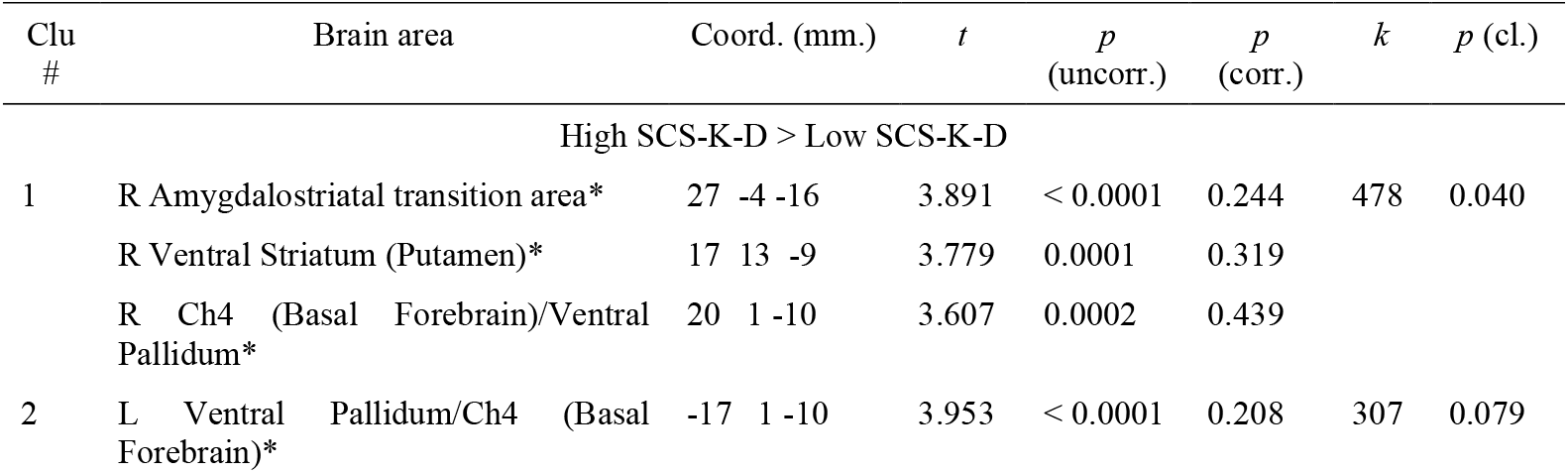
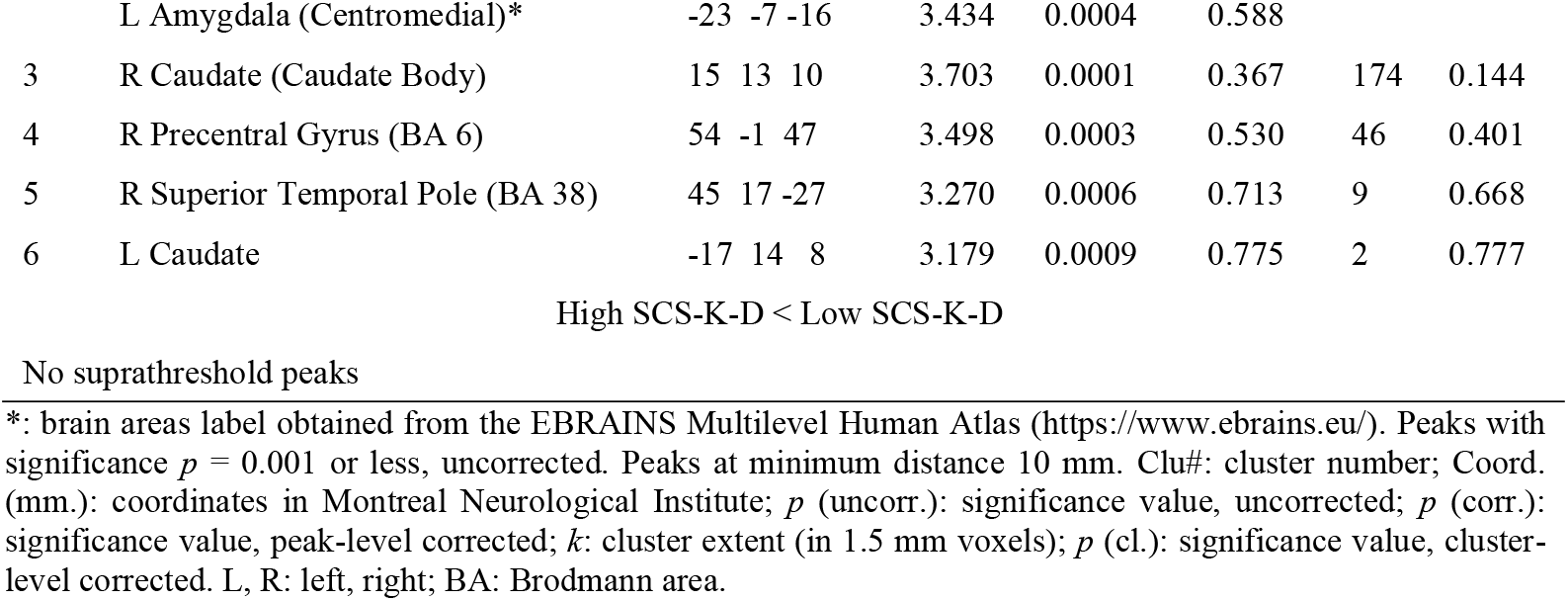
Whole brain effects of self-regulation on time on task.

### 3.3 Association of self-regulation with the interaction between time on task and reward (hypothesis 3)

Relatively poorly regulated individuals showed increased VTA/SN recruitment when considering the interaction between time on task and reward (see Table 3A in the Appendix, High SCS-K-D < Low SCS-K-D, cluster #6), consistent with hypothesis 3 (Figure 3A). However, the result did not survive stringent correction for multiple testing, reaching significance only at trend level (x, y, z: -8, -16, -15, *t* = -2.441, *p* = 0.072). This result was driven by low reward trials. While in these trials the activity differed between individuals according to their self-regulation score, with highly regulated individuals showing the positive trend generally found in this area (*t* = 2.003), in high reward trials there were no individual differences from self-regulation (*t* = -0.813). This finding is consistent with highly regulated individuals having a similar signal in low and high reward trials (Figure 3B).

**Table 3A.**
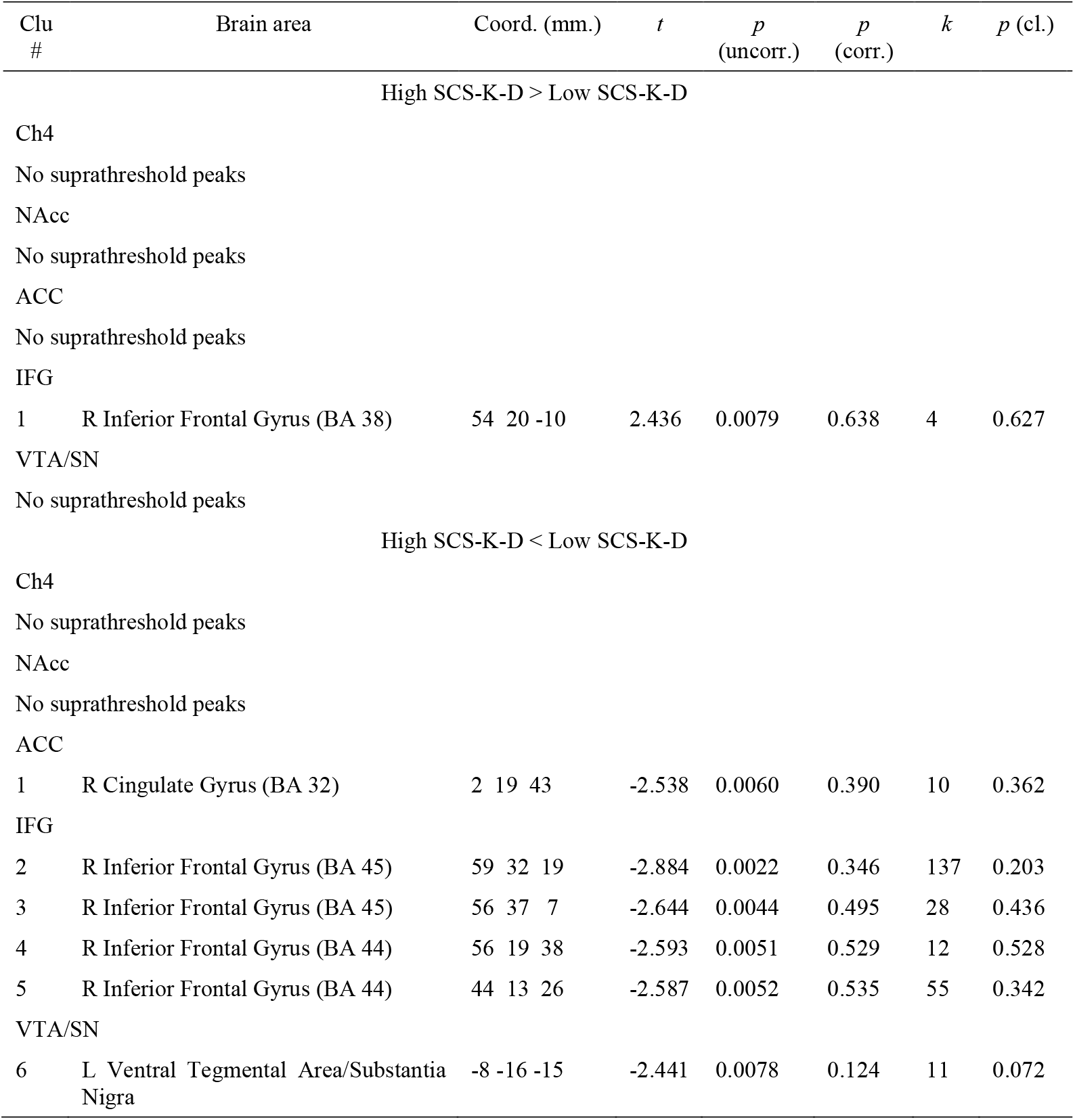
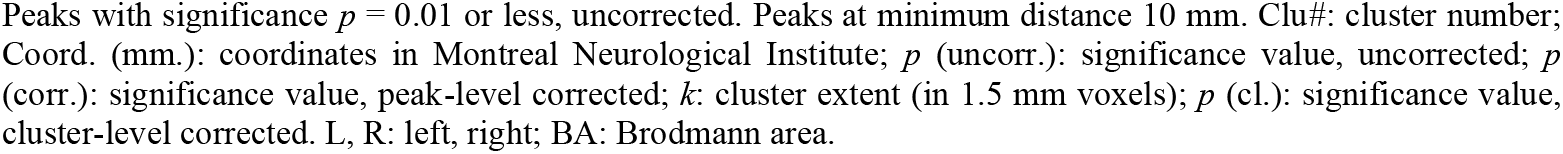
ROI effects of self-regulation on the interaction between time on task and reward.

| Clu # | Brain area | Coord. (mm.) | $t$ | $p$ (uncorr.) | $p$ (corr.) | $k$ | $p$ (cl.) |
| --- | --- | --- | --- | --- | --- | --- | --- |
| High SCS-K-D > Low SCS-K-D |  |  |  |  |  |  |  |
| Ch4 |  |  |  |  |  |  |  |
| No suprathreshold peaks |  |  |  |  |  |  |  |
| NAcc |  |  |  |  |  |  |  |
| No suprathreshold peaks |  |  |  |  |  |  |  |
| ACC |  |  |  |  |  |  |  |
| No suprathreshold peaks |  |  |  |  |  |  |  |
| IFG |  |  |  |  |  |  |  |
| 1 | R Inferior Frontal Gyrus (BA 38) | 54 20 -10 | 2.436 | 0.0079 | 0.638 | 4 | 0.627 |
| VTA/SN |  |  |  |  |  |  |  |
| No suprathreshold peaks |  |  |  |  |  |  |  |
| High SCS-K-D < Low SCS-K-D |  |  |  |  |  |  |  |
| Ch4 |  |  |  |  |  |  |  |
| No suprathreshold peaks |  |  |  |  |  |  |  |
| NAcc |  |  |  |  |  |  |  |
| No suprathreshold peaks |  |  |  |  |  |  |  |
| ACC |  |  |  |  |  |  |  |
| 1 | R Cingulate Gyrus (BA 32) | 2 19 43 | -2.538 | 0.0060 | 0.390 | 10 | 0.362 |
| IFG |  |  |  |  |  |  |  |
| 2 | R Inferior Frontal Gyrus (BA 45) | 59 32 19 | -2.884 | 0.0022 | 0.346 | 137 | 0.203 |
| 3 | R Inferior Frontal Gyrus (BA 45) | 56 37 7 | -2.644 | 0.0044 | 0.495 | 28 | 0.436 |
| 4 | R Inferior Frontal Gyrus (BA 44) | 56 19 38 | -2.593 | 0.0051 | 0.529 | 12 | 0.528 |
| 5 | R Inferior Frontal Gyrus (BA 44) | 44 13 26 | -2.587 | 0.0052 | 0.535 | 55 | 0.342 |
| VTA/SN |  |  |  |  |  |  |  |
| 6 | L Ventral Tegmental Area/Substantia Nigra | -8 -16 -15 | -2.441 | 0.0078 | 0.124 | 11 | 0.072 |

Peaks with significance $p = 0.01$ or less, uncorrected. Peaks at minimum distance 10 mm. Clu#: cluster number; Coord. (mm.): coordinates in Montreal Neurological Institute; $p$ (uncorr.): significance value, uncorrected; $p$ (corr.): significance value, peak-level corrected; $k$ : cluster extent (in 1.5 mm voxels); $p$ (cl.): significance value, cluster-level corrected. L, R: left, right; BA: Brodmann area.

**Figure 3.**
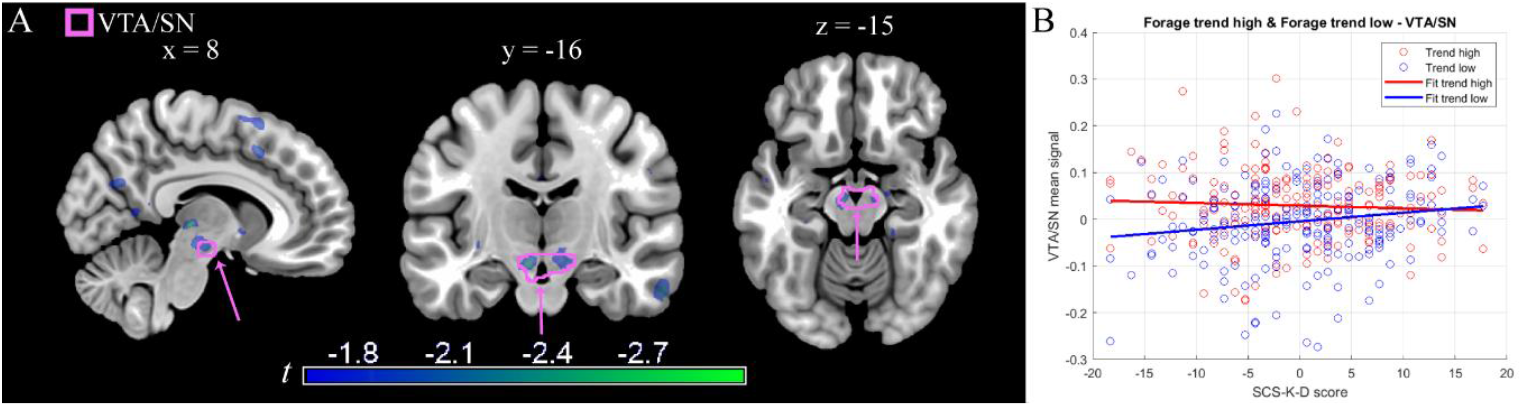
A: Whole brain functional activity of the interaction between time on task and reward levels × self-regulation scores. The VTA/SN ROI mask (in pink) is superimposed on the *t* map (thresholded for illustration purposes at -3 < *t* < -1.65, *p* < 0.05 uncorrected). The peak in the posterior thalamus visible in the transversal slice at x = 8 is bilateral and located in correspondence of the habenula, but failed to survive correction (Table 4A, cluster #3). B: Illustrative plot of the time on task signal on self-regulation levels for high (red) and low (blue) reward trials in the voxels of the interaction, showing differences between participants at low reward levels (see text for inference).

The whole brain analysis found no evidence of a significant effect of self-regulation traits in the interaction between time on task and reward (Table 4A, Appendix).

**Table 4A.** Whole brain effects of self-regulation on the interaction between time on task and reward.

| Clu<br># | Brain area | Coord. (mm.) | $t$ | $p$<br>(uncorr.) | $p$<br>(corr.) | $k$ | $p$ (cl.) |
| --- | --- | --- | --- | --- | --- | --- | --- |
| High SCS-K-D > Low SCS-K-D |  |  |  |  |  |  |  |
| No suprathreshold peaks |  |  |  |  |  |  |  |
| High SCS-K-D < Low SCS-K-D |  |  |  |  |  |  |  |
| 1 | L Superior Frontal Gyrus (BA 6) | -23 10 59 | -3.502 | 0.0003 | 0.611 | 41 | 0.506 |
| 2 | R Superior Frontal Gyrus (BA 8) | 29 10 58 | -3.239 | 0.0007 | 0.815 | 37 | 0.529 |
| 3 | R/L Habenular Nucleus | 3 -24 2 | -3.372 | 0.0004 | 0.717 | 9 | 0.758 |
| 4 | R Angular Gyrus (BA 39) | 42 -64 34 | -3.188 | 0.0008 | 0.846 | 1 | 0.877 |
| 5 | L Supplementary Motor Area (BA 6) | -18 2 64 | -3.162 | 0.0009 | 0.861 | 1 | 0.877 |

Peaks with significance $p = 0.001$ or less, uncorrected. Peaks at minimum distance 10 mm. Clu#: cluster number; Coord. (mm.): coordinates in Montreal Neurological Institute; $p$ (uncorr.): significance value, uncorrected; $p$ (corr.): significance value, peak-level corrected; $k$ : cluster extent (in 1.5 mm voxels); $p$ (cl.): significance value, cluster-level corrected. L, R: left, right; BA: Brodmann area.

## 4 Discussion

### 4.1 Behavioral data

The lack of associations between self-regulation levels and performance decrements is consistent with previous findings in the human literature for reaction times (Becker et al., 2015; Harwood et al., 2023) and accuracy (Becker et al., 2015).

### 4.2 Time on task

The involvement of frontal and prefrontal areas in the implementation of self-control (Banfield et al., 2004; Heatherton & Wagner, 2011; Posner et al., 2007), as predicted by the first hypothesis, found no confirmation in the present study. Even at a ROI level, there was no recruitment of the ACC or the IFG. Beyond their involvement in self-control (Banfield et al., 2004; Casey et al., 2011; Posner et al., 2007), ACC and IFG were among the most active areas in this dataset in the model of time on task and its interaction with reward levels (Orsini et al., 2026), demonstrating the capacity of our paradigm to recruit these substrates.

In models from the neuroimaging literature on cognitive effort, ACC activity is thought to be related to the computation of the amount and type of effort needed to maximize the difference between expected profits and intrinsic costs of effort (Shenhav et al., 2013; Shenhav et al., 2016; Shenhav et al., 2017). Therefore, the null finding arising from our paradigm should be interpreted in this more restrictive context, suggesting that self-regulation may not be associated with differences in activity of substrates of effort evaluation or its computation. However, cognitive effort evaluation and working memory load might be distinct processes. In this case, our task may not have been the appropriate approach to elicit the phenotype of cognitive control that was at the basis of our first hypothesis. Indeed, no working memory is required to execute our task, which is meant to capture neural correlates of the effort to remain on task rather than cognitive load *per se*.

Another possible issue with this null finding is that our assessment of self-regulation may not match the notion of cognitive control as operationalized elsewhere. It should be noted, however, that in our method development we verified that our self-regulation scores were associated with behavioral indices of self-regulation reported in the literature, such as lower BMI (see Rating scales section in Material and methods). This argues in favor of our measurement of self-regulation as capturing a meaningful aspect of this broad construct.

In contrast, the association of the SCS scores of our study with the activity in NBM, a structure known to be crucial in sustaining attention (Harati et al., 2008; Liu et al., 2018; Martinez & Sarter, 2004; McGaughy et al., 2002; McGaughy et al., 1999), was consistent with our second hypothesis. During the cognitive effort induced by time on task, highly regulated individuals exhibited a greater functional involvement of the NBM in the portion consistent with cholinergic activity (Ch4). This finding is further supported by the fact that the inference about BF involvement did not depend on ROI analyses. The whole-brain analysis also suggested a more extensive involvement of this brain region, including substrates spanning from the ventral pallidum to the centromedial and transitional parts of the amygdala, where NBM cells intermingle (Mesulam, 2013). Congruently with this evidence, a previous study on individual differences in impulsivity identified the presence of a negative correlation between impulsivity and dorsal amygdala activity (Brown et al., 2006), peaking in a region that we view as compatible with recruitment of NBM/Ch4.

Studies in laboratory animals have provided evidence on the cholinergic BF as part of a network responsible for increasing ACh cortical efflux, mediating cognitive effort during cognitive challenges in sustained attention tasks (Sarter et al., 2006). This framework suggests a role of cholinergic activity in the interaction between the time spent on task (a cognitive challenge) and trait self-regulation detected in the present study. This result is also consistent with evidence from the animal literature on the relationship between the cholinergic system and individual differences during effort exertion (Hosking et al., 2014). The phenotype-dependent variations in Ch4 activity observed in the current study might be related to differences in the amount of ACh efflux towards the cortex, with consequences on cognitive effort exertion. Future studies combining functional neuroimaging and ACh recording or ACh agents’ administration will be needed to clarify these effects.

### 4.3 Interaction between time on task and reward

Subcortically, the VTA/SN activity evoked by the time spent on task and by its interaction with reward levels (Orsini et al., 2026) did not reach the pre-established threshold for statistical significance with the significance correction method. This analysis only yielded a nominally significant association in the VTA/SN. While we acknowledge that findings failing to meet correction thresholds must be interpreted with caution, we elected to report this pattern for several reasons. First, its selective anatomical distribution corresponds to substrates of reward processing, as hypothesized. Second, it aligned with our third hypothesis on the importance of substrates associated with reward detection on the sensitivity to reward of less regulated individuals. In our task, participants were required to reach a minimum score of corrected responses to receive the reward accumulated during the experiment, forcing them to respond to low as well as high reward trials to reach this threshold. In our data, participants with high self-regulation scores showed no differences in the time on task signal at different levels of reward, while this signal was lower in low reward trials in low self-regulation participants.

Previous studies have reported the dopaminergic (in particular D2 and D3) autoreceptor availability in the VTA and SN to be negatively correlated with trait impulsivity in humans (Buckholtz et al., 2010). Other studies in laboratory animals have shown a modulation of impulsive behavior and attention after the manipulation of VTA neurons and VTA-NAcc projections (Boekhoudt et al., 2017; Flores-Dourojeanni et al., 2023; Flores-Dourojeanni et al., 2021). However, there are relatively few studies focusing on VTA/SN and several aspects of its sensitivity to dopamine activity remain to be understood. These findings argue for the need to better clarify the possible VTA or dopaminergic involvement in self-regulation-related processes and impulsivity.

## 5 Conclusion

The present findings suggest that self-regulation in humans may operate through multiple dissociable neural mechanisms. The behavioral literature has primarily linked self-regulation to reward-related decision making, particularly delay discounting. Our findings in the VTA/SN—showing differential modulation by reward in less regulated individuals—are consistent with this established framework.

The current study presents evidence of a second and potentially independent mechanism. Highly regulated individuals present a stronger involvement of Ch4 compared to their less regulated counterparts over the time on task, independently of reward levels. This evidence supports the existence of an association between the cholinergic system and self-regulation providing information on the possible mechanism through which self-regulation affects cognition, suggesting it may support a form of attentional maintenance that has remained behaviorally undetected in standard paradigms. Future studies integrating cognitive testing, neuroimaging and neuropharmacology will be needed to better clarify the association between the cholinergic system and the ACh efflux and measures of self-regulation.

We emphasize that a complex trait such as self-regulation in humans is not likely to be explained by a single intermediate phenotype. Cholinergic activity may represent a fundamental dimension of self-regulation that complements the well-established reward-sensitivity dimension.

## Data and Code Availability

Data and code may be shared under a formal data sharing agreement which includes the purposes of the analyses of the requesting researchers. These purposes will need to be consistent with those to which participants agreed at data collection, as stated in the written consent. The study will need to be approved by the requesting researcher’s ethics committee.

## Contributions

Designed research: RV, CO. Funding: RV. Collected data: JEB. Wrote software: RV, DAH. Analyzed data: CO. Wrote manuscript: CO, RV, KL. All authors revised and approved the final version of the manuscript.

## Funding, Declaration of Competing Interest and Acknowledgements

A preliminary version of this work was presented at the CogBases workshop (Paris, Institut Pasteur, 10-11 October 2023). This work was supported in whole by an ERA-PERMED grant (project ArtiPro) of the FWF Austrian Science Fund (grant number I 5903) [Grant-DOI: 10.55776/I5903] to Roberto Viviani. The authors declare no conflict of interest. For open access purposes, the author has applied a CC BY public copyright license to any manuscript version arising from this submission.

## Appendix

**Table 1A.**
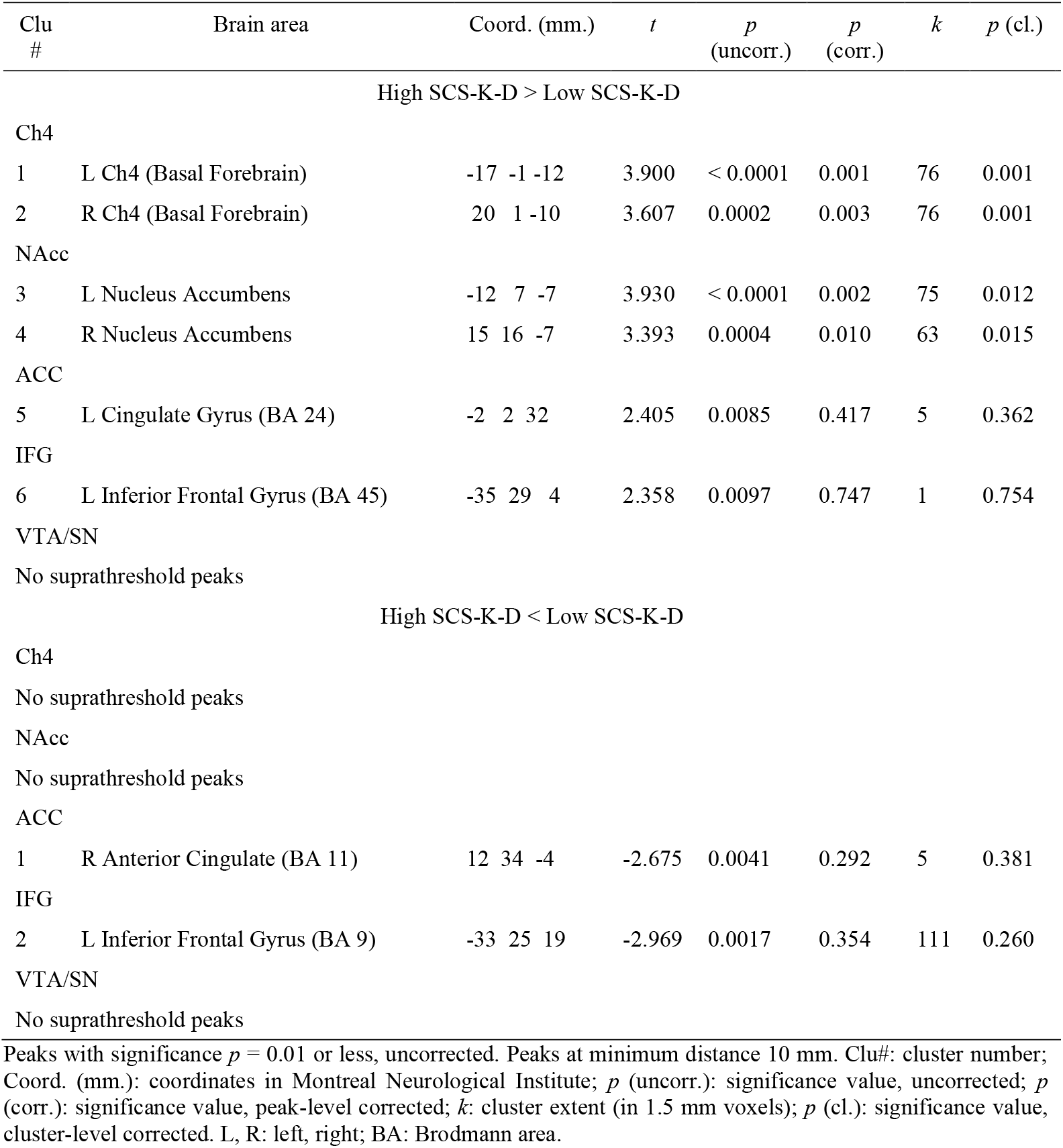
ROI effects of self-regulation on time on task.

