## Supplementary Materials for "Basal forebrain and neural correlates of self-regulation traits in sustained attention"

### Supplementary R Markdown

#### Load data

```
data203 <- read.table("Query_Foraging_behavioral_203part.txt", header = T)
```

Information about the interstimulus interval was calculated.

```
RTData <- read.table('RTData.txt', header = T,  sep='\t')
RTData <- RTData[ RTData$SubjectID %in% data203$SubjectID, ]

index <- RTData$ISI > 0
summary(RTData$ISI[index])
```

```
##    Min. 1st Qu.  Median    Mean 3rd Qu.    Max.
##  0.7995  1.0133  1.2133  1.2284  1.4134  1.8181
```

#### Descriptive

We visualized the distribution of our main predictor variable, self-regulation, to verify the absence of extreme values that may constitute high leverage observations.

```
hist(data203$SCS_K, xlab = "SCS-K-D scores", main = "Self-regulation Histogram")
```

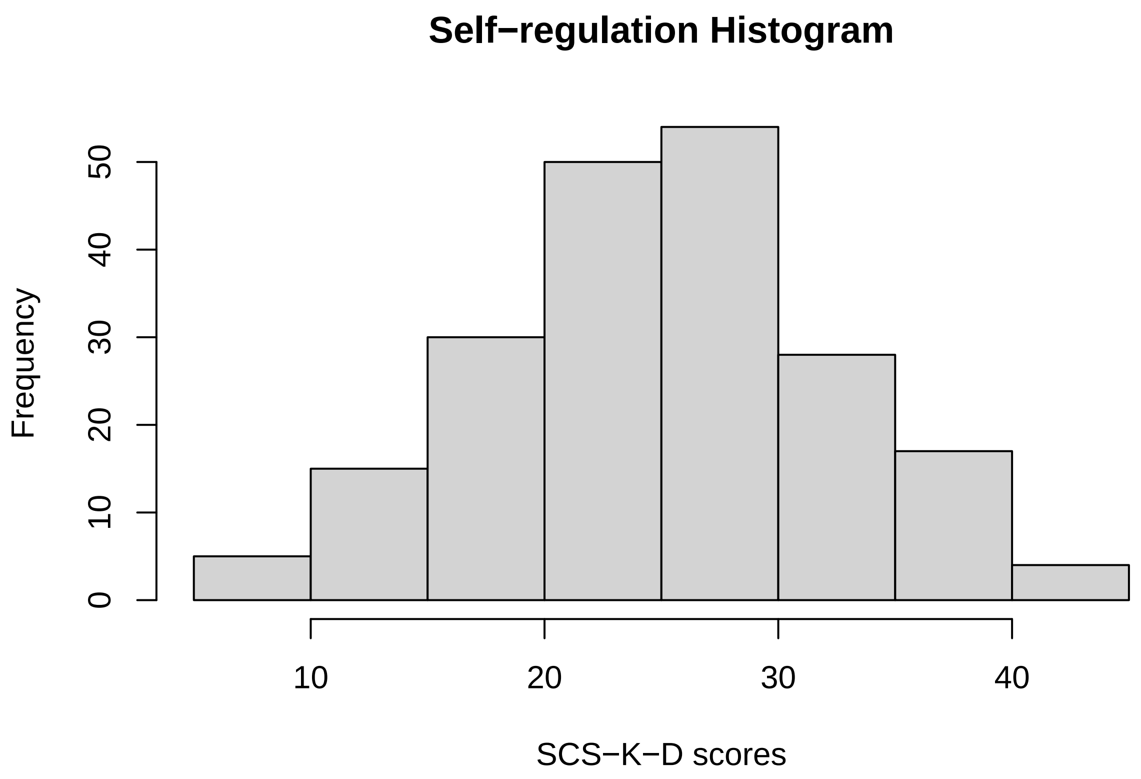


The distribution appears to be roughly normal and does not show any outlying value.

We obtained demographic information like the number of females and participants from Bonn, and the mean age to characterize the sample in the manuscript.

```
sum(data203$Bonn == 1)
```

```
## [1] 20
```

```
sum(data203$Female == 1)
```

```
## [1] 114
```

```
summary(data203$Age)
```

```
##    Min. 1st Qu.  Median    Mean 3rd Qu.    Max.
##   18.00   20.00   22.00   23.06   25.00   38.00
```

```
sd(data203$Age)
```

```
## [1] 3.881468
```

### Modeling

#### Self-regulation and demographics

We characterized our self-regulation measure in terms of the demographics of our sample.

```r
fit1_203 <- lm(SCS_K_D ~ Bonn + Age + Female, data=data203)
summary(fit1_203)
```

```
##
## Call:
## lm(formula = SCS_K_D ~ Bonn + Age + Female, data = data203)
##
## Residuals:
##      Min       1Q   Median       3Q      Max
## -19.5006  -5.1832  -0.2224   5.0472  19.4271
##
## Coefficients:
##             Estimate Std. Error t value Pr(>|t|)
## (Intercept)  21.4292     3.4441   6.222 2.84e-09 ***
## Bonn         -4.1816     1.9154  -2.183   0.0302 *
## Age           0.1270     0.1475   0.861   0.3905
## Female        2.4051     1.0527   2.285   0.0234 *
## ---
## Signif. codes:  0 '***' 0.001 '**' 0.01 '*' 0.05 '.' 0.1 ' ' 1
##
## Residual standard error: 7.437 on 199 degrees of freedom
## Multiple R-squared:  0.04746,    Adjusted R-squared:  0.0331
## F-statistic: 3.305 on 3 and 199 DF,  p-value: 0.02129
```

Females were significantly associated with a higher self-regulation score, while age showed no association.

#### Self-regulation and body mass index (BMI)

We tested the association between the self-regulation score and BMI, which was available in our database. Test had the purpose to demonstrate the validity of the rating scale we used, given the reports in the literature on this association.

```r
fitBMI_203 <- lm(BMI ~ SCS_K_D + Bonn + Age + Female, data=data203)
summary(fitBMI_203)
```

```
##
## Call:
## lm(formula = BMI ~ SCS_K_D + Bonn + Age + Female, data = data203)
##
## Residuals:
##     Min      1Q  Median      3Q     Max
## -6.2467 -1.9279 -0.4265  1.6611 11.9390
##
## Coefficients:
##             Estimate Std. Error t value Pr(>|t|)
## (Intercept) 21.20246    1.54486  13.724   <2e-16 ***
```

```
## SCS_K_D     -0.07308    0.02909  -2.512   0.0128 *
## Bonn         0.38280    0.79547   0.481   0.6309
## Age          0.17179    0.06066   2.832   0.0051 **
## Female      -0.79834    0.43767  -1.824   0.0697 .
## ---
## Signif. codes:  0 '***' 0.001 '**' 0.01 '*' 0.05 '.' 0.1 ' ' 1
##
## Residual standard error: 3.052 on 198 degrees of freedom
## Multiple R-squared:  0.1071, Adjusted R-squared:  0.08909
## F-statistic: 5.939 on 4 and 198 DF,  p-value: 0.0001559
```

As reported in the literature, self-regulation was a significant negative predictor of BMI, beside the positive association with age. This supports the validity of the self-regulation scale we used in the experiment.

#### Self-regulation and performance (RTs and Accuracy)

Task-related effects on the behavioral performance are reported in Orsini et al., 2026 on a larger sample (415 participants) that included the participants from the present study (203 participants). Here, we investigate on presence of an association between trait-self regulation and RTs and Accuracy on those participants who filled out the Self-Control Scale.

First, to support lmer convergence, a few variables are scaled. The inter-stimulus interval (ISI) and the switch are centered so to have a neutral effect in the first trials in block. These trials, indeed, have no switch or ISI.

```r
SummaryData <- read.table('SummaryData.txt', header = T)
SummaryData <- SummaryData[ SummaryData$SubjectID %in% data203$SubjectID, ]
RTData <- rename(RTData, 'switch' = 'PI')

data203_RTs <- left_join(RTData, data203, by = "SubjectID") %>% left_join(SummaryData,
by = "SubjectID")

data203_RTs$block <- data203_RTs$block / 10
data203_RTs$firstTrialInBlock <- as.numeric(data203_RTs$trialInBlock == 1)
data203_RTs$trialInBlock <- data203_RTs$trialInBlock / 10

data203_RTs$ISI[index] <- data203_RTs$ISI[index] - mean(data203_RTs$ISI[index])
data203_RTs$switch[index] <- data203_RTs$switch[index] -
mean(data203_RTs$switch[index])
sum(is.na(data203_RTs))
```

```
## [1] 90
```

As in Orsini et al., 2026, RTs were thresholded ($0.25 < RTs < 0.8$) to avoid excessively rapid and slow responses, and an Accuracy dataframe is generated.

```r
ggplot(data203_RTs, aes(x= trial, y=RT)) + geom_point(alpha = 0.1) + labs(title =
"RTs", x = "Trials", y = "RTs (sec)") + geom_hline(yintercept = 0.25, color = "red") +
geom_hline(yintercept = 0.8, color = "blue") + theme_minimal()
```

```
## Warning: Removed 90 rows containing missing values or values outside the scale range
## (`geom_point()`).
```

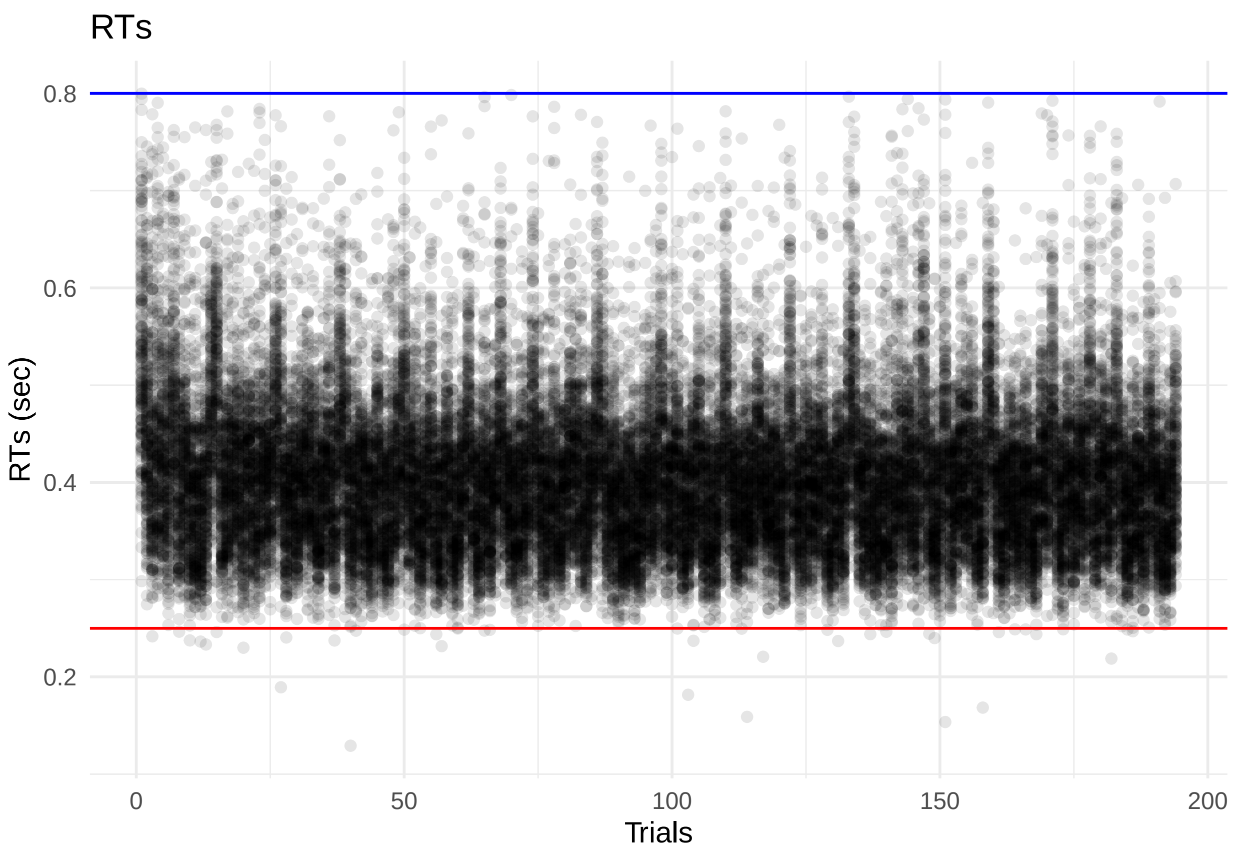


```
data203_Accuracy <- filter(data203_RTs, response == "hit" | response == "incorrect" |
    response == "miss")

data203_RTs <- filter(data203_RTs, response == "hit" | response == "incorrect", RT > 0.25,
    RT < 0.8)
```

**RTs** We check on the presence of an association between SCS scores and RTs.

```
data203_RTs$ISIms <- data203_RTs$ISI * 1000
summary(lmer(RT*1000 ~ firstTrialInBlock + trialInBlock + SCS_K_D + switch + ISIms +
             I(code == "high") + block + Bonn + Age + Female + (1 | SubjectID),
             data = data203_RTs))
```

```
## Linear mixed model fit by REML. t-tests use Satterthwaite's method [
## lmerModLmerTest]
## Formula: RT * 1000 ~ firstTrialInBlock + trialInBlock + SCS_K_D + switch +
##     ISIms + I(code == "high") + block + Bonn + Age + Female +
##     (1 | SubjectID)
##    Data: data203_RTs
##
## REML criterion at convergence: 419519.6
##
## Scaled residuals:
##     Min      1Q  Median      3Q     Max
```

```
## -3.9505 -0.6045 -0.1256  0.4297  7.2932
##
## Random effects:
##  Groups    Name        Variance Std.Dev.
##  SubjectID (Intercept) 2845     53.34
##  Residual              2501     50.01
## Number of obs: 39246, groups:  SubjectID, 203
##
## Fixed effects:
##                          Estimate Std. Error         df t value Pr(>|t|)
## (Intercept)             3.694e+02  2.707e+01  1.983e+02  13.646  < 2e-16 ***
## firstTrialInBlock       8.683e+01  1.048e+00  3.904e+04  82.870  < 2e-16 ***
## trialInBlock            1.443e+00  8.281e-01  3.904e+04   1.742   0.0815 .
## SCS_K_D                 4.336e-01  5.096e-01  1.980e+02   0.851   0.3958
## switch                 -2.729e+01  5.361e-01  3.904e+04 -50.900  < 2e-16 ***
## ISIms                  -5.704e-02  9.584e-04  3.904e+04 -59.520  < 2e-16 ***
## I(code == "high")TRUE -3.599e+00  5.075e-01  3.904e+04  -7.091 1.36e-12 ***
## block                  -1.171e+01  5.486e-01  3.904e+04 -21.344  < 2e-16 ***
## Bonn                   -6.146e+01  1.393e+01  1.980e+02  -4.411 1.69e-05 ***
## Age                     1.570e+00  1.063e+00  1.980e+02   1.477   0.1412
## Female                  5.059e+00  7.666e+00  1.980e+02   0.660   0.5101
## ---
## Signif. codes:  0 '***' 0.001 '**' 0.01 '*' 0.05 '.' 0.1 ' ' 1
##
## Correlation of Fixed Effects:
##             (Intr) frsTIB trlInB SCS_K_ switch ISIms  I(=="" block  Bonn
## frstTrlInBl -0.013
## trialInBlck -0.022  0.480
## SCS_K_D     -0.403  0.000  0.000
## switch      -0.001  0.021  0.043  0.000
## ISIms        0.002 -0.053 -0.111  0.000  0.088
## I(=="")TRUE -0.008 -0.001 -0.003  0.000 -0.046 -0.007
## block       -0.017  0.004  0.009  0.000  0.018  0.007 -0.093
## Bonn         0.251  0.000  0.000  0.153  0.000  0.000  0.000  0.000
## Age         -0.863  0.000  0.000 -0.061  0.000  0.000  0.000  0.000 -0.409
## Female      -0.124  0.000  0.000 -0.160  0.000  0.000  0.000  0.000 -0.033
##             Age
## frstTrlInBl
## trialInBlck
## SCS_K_D
## switch
## ISIms
## I(=="")TRUE
## block
## Bonn
## Age
## Female       0.047
```

The model shows that the variable SCS_K_D has no significant effect on the RTs.

**Accuracy** We analyse Accuracy. Global accuracy is over the 99%.

```r
summary(data203_Accuracy$response) #39369 total hit/miss/incorrect responses
```

```
##    Length     Class      Mode 
##     39369 character character
```

```r
summary(data203_Accuracy$response == "hit")
```

```
##    Mode   FALSE    TRUE 
## logical     186   39183
```

```r
accuracy_percent <- (39183/39369)*100
accuracy_percent
```

```
## [1] 99.52755
```

We center some predictors to aid convergence.

```r
data203_Accuracy$firstTrialInBlock_centered <- scale(data203_Accuracy$firstTrialInBlock,
                                                     center = T, scale = F)
data203_Accuracy$block_centered <- scale(data203_Accuracy$block, center = T, scale = F)
data203_Accuracy$trialInBlock_centered <- scale(data203_Accuracy$trialInBlock, center = T,
                                                scale = F)
data203_Accuracy$SCS_K_D_centered <- scale(data203_Accuracy$SCS_K_D, center = T,
                                           scale = F)
data203_Accuracy$Age_centered <- scale(data203_Accuracy$Age, center = T, scale = F)
```

The analysis of accuracy does not show an effect of SCS scores on Accuracy.

```r
summary((glmer(response == "hit" ~ firstTrialInBlock_centered +
    trialInBlock_centered + SCS_K_D_centered + Age_centered + Female + switch + ISI +
      I(code == "high") + block_centered + Bonn + (1 | SubjectID),
    data = data203_Accuracy, family = binomial, control = glmerControl(optimizer =
    "bobyqa", optCtrl = list(maxfun = 100000)))))
```

```
## Generalized linear mixed model fit by maximum likelihood (Laplace
##   Approximation) [glmerMod]
##  Family: binomial  ( logit )
## Formula: 
## response == "hit" ~ firstTrialInBlock_centered + trialInBlock_centered +  
##     SCS_K_D_centered + Age_centered + Female + switch + ISI +  
##     I(code == "high") + block_centered + Bonn + (1 | SubjectID)
##    Data: data203_Accuracy
## Control: glmerControl(optimizer = "bobyqa", optCtrl = list(maxfun = 1e+05))
## 
##      AIC      BIC   logLik deviance df.resid 
##   2222.3   2325.2  -1099.1   2198.3    39357 
## 
## Scaled residuals: 
##     Min      1Q  Median      3Q     Max 
## -36.807   0.032   0.050   0.071   0.364
```

```
##
## Random effects:
##  Groups    Name        Variance Std.Dev.
##  SubjectID (Intercept) 0.808    0.8989
## Number of obs: 39369, groups:  SubjectID, 203
##
## Fixed effects:
##                            Estimate Std. Error z value Pr(>|z|)
## (Intercept)                 5.864598   0.200431  29.260  < 2e-16 ***
## firstTrialInBlock_centered -1.500123   0.283255  -5.296 1.18e-07 ***
## trialInBlock_centered      -0.876912   0.257355  -3.407 0.000656 ***
## SCS_K_D_centered           -0.009023   0.013967  -0.646 0.518284
## Age_centered               -0.029114   0.028464  -1.023 0.306382
## Female                     -0.006986   0.212822  -0.033 0.973814
## switch                      1.518924   0.191319   7.939 2.03e-15 ***
## ISI                         0.570298   0.295809   1.928 0.053864 .
## I(code == "high")TRUE       0.418701   0.151782   2.759 0.005806 **
## block_centered             -0.030183   0.160191  -0.188 0.850546
## Bonn                        0.324307   0.397696   0.815 0.414807
## ---
## Signif. codes:  0 '***' 0.001 '**' 0.01 '*' 0.05 '.' 0.1 ' ' 1
##
## Correlation of Fixed Effects:
##             (Intr) frTIB_ trlIB_ SCS_K_ Ag_cnt Female switch ISI    I(==""
## frstTrlInB_ -0.181
## trlInBlck_c -0.096  0.629
## SCS_K_D_cnt  0.078  0.001  0.001
## Age_centerd  0.030  0.001  0.001 -0.052
## Female      -0.585  0.000  0.000 -0.178  0.043
## switch       0.313 -0.212  0.032  0.000 -0.001  0.000
## ISI          0.058 -0.089 -0.085  0.000  0.000  0.000  0.082
## I(=="")TRUE -0.299 -0.015 -0.023  0.000  0.000  0.000 -0.033 -0.004
## block_cntrd  0.002 -0.006 -0.002  0.000  0.000  0.000  0.052  0.072  0.011
## Bonn        -0.175  0.000  0.000  0.157 -0.370 -0.020  0.001  0.001  0.000
##             blck_c
## frstTrlInB_
## trlInBlck_c
## SCS_K_D_cnt
## Age_centerd
## Female
## switch
## ISI
## I(=="")TRUE
## block_cntrd
## Bonn         0.000
```
